# Structure-guided design of novel orthosteric inhibitors of integrin αIIbβ3 that prevent thrombosis but preserve hemostasis

**DOI:** 10.1101/509299

**Authors:** Brian D. Adair, José L. Alonso, Johannes van Agthoven, Vincent Hayes, Hyun Sook Ahn, I-Shing Yu, Shu-Wha Lin, Mortimer Poncz, Jian-Ping Xiong, M. Amin Arnaout

## Abstract

A prevailing dogma is that effective inhibition of vascular thrombosis by therapeutic targeting of platelet integrin αIIbβ3 cannot be achieved without compromising hemostasis. The resulting serious bleeding and increased morbidity and mortality have limited use of current anti-αIIbβ3 drugs to high-risk cardiac patients. It is speculated that these adverse outcomes result from drug-induced conformational changes in αIIbβ3 but direct proof is lacking. We used structure-guided design to generate the ligand-mimetic peptide Hr10 and a modified form of the partial agonist drug Tirofiban that now act as “pure” orthosteric antagonists of αIIbβ3, i.e. they no longer induce the conformational changes in αIIbβ3 and also suppress these changes in presence of agonists. Both agents inhibited human platelet aggregation effectively but without interfering with clot retraction. When tested in a humanized mouse model of thrombosis predictive of clinical efficacy, Hr10 was as effective as the αIIbβ3 partial agonist peptide drug Eptifibatide in inhibiting arteriolar thrombosis, but in sharp contrast to Eptifibatide, Hr10 did not cause serious bleeding, establishing a causal link between partial agonism and impaired hemostasis. Pure orthosteric inhibitors of αIIbβ3 may thus offer safer alternatives for human therapy. Our structure-guided approach may also find utility in designing similar drug candidates targeting other integrins and in providing vital tools for further probing structure-activity relationships in integrins.

## Introduction

Platelet activation and accumulation at the site of blood vessel injury are the initial steps in hemostasis. When activated by several agonists including adenosine diphosphate (ADP), thrombin or collagen, platelets adhere to the disrupted surface, and aggregate upon binding of soluble fibrinogen in circulating blood to agonist-activated αIIbβ3 ^1^. Fibrin generated by thrombin at or near the platelet surface also binds αIIbβ3, driving clot retraction ^2^, thereby consolidating the integrity of the hemostatic plug, restoring blood flow and promoting wound closure ^3^. Excessive platelet activation by agonists may lead to formation of occlusive thrombi, which are responsible for acute myocardial infarction and stroke ^4^, hemodialysis access failure ^5^, early loss of kidney allograft ^6^, tumor growth and metastasis ^7^, and may also contribute to fibril formation in cerebral vessels of Alzheimer's disease patients ^8^.

The three parenteral anti-αIIbβ3 drugs Eptifibatide, Tirofiban and Abciximab (which additionally inhibits αVβ3) have demonstrated efficacy in reducing death and ischemic complications in victims of heart attacks ^9^. However, their clinical use in acute coronary syndrome has been associated with serious bleeding, which often requires cessation of therapy, putting heart attack victims at high risk of re-thrombosis. And orally active anti-αIIbβ3 agents given to patients at risk of acute coronary syndromes were abandoned because of increased risk of patient death linked to paradoxical coronary thrombosis ^10–12^. Concluding that the adverse outcomes resulting from targeting αIIbβ3 are unavoidable, pharmaceutical companies developed inhibitors of the platelet ADP receptor P_2_Y_12_ and thrombin receptor PAR1, both upstream of αIIbβ3. However, a considerable number of patients receiving these newer drugs continue to experience serious bleeding and thrombotic events ^13–15^. Thus, there remains an unmet clinical need for new anti-thrombosis drugs that maintain efficacy while preserving hemostasis ^16^.

Previous studies have shown that the three current anti-αIIbβ3 drugs are partial agonists, i.e. they trigger large activating conformational changes in αIIbβ3 that enhance receptor binding to physiologic ligand, promoting thrombosis ^17,18^, and also expose neo-epitopes for natural antibodies, causing immune thrombocytopenia ^19^. X-ray structures of unliganded and ligand-bound integrins ^20–22^ showed that the conformational changes in the integrin induced by binding of ligands or ligand-mimetic drugs are initiated in the integrin ligand-binding vWFA domain (A-domain). Ligand binding to β3 integrins triggers tertiary changes in the A-domain of the β3 subunit (βA domain), comprising the inward movement of the N-terminal α1 helix (reported by movement of β3-Tyr^122^) towards the Mg^2+^ or Mn^2+^ ion coordinated at the *m*etal *i*on-*d*ependent *a*dhesion *s*ite (MIDAS). This reshapes the C-terminal F-α7 loop and repositions the α7 helix causing a swing-out of the hybrid domain underneath the βA domain. This movement converts the integrin from the genu-bent to the extended conformation, separates the transmembrane and cytoplasmic tails that allows formation of integrin-cytoskeleton interactions mediating dynamic cell adhesion ^23^.

Recently, we have shown that the above conformational changes in integrin αVβ3 can be prevented by binding of the ligand hFN10, a mutant form of the 10^th^ type III domain of fibronectin (FN10) ^24^. This unexpected effect was traced to a key π-π interaction between the indole derivative, tryptophan, that immediately follows the RGD motif of hFN10, and the β3-Tyr^122^ in the ligand binding βA domain of αVβ3 ^24^. The applicability of this structural feature for generating novel antagonists for other integrins has not been explored. In this manuscript we used this information to engineer pure antagonists of αIIbβ3 that are effective in preventing arteriolar thrombosis while preserving hemostasis, thus establishing a causal link between the drug-induced conformational changes in integrins and adverse outcomes, and paving the way for potentially safer medical therapies.

## Results

### Development of peptide Hr10

The inability of hFN10 to bind αIIbβ3 ^25^ was investigated by superimposing the βA domains from the crystal structures of αIIbβ3/Eptifibatide complex (2vdn.pdb) and αVβ3/hFN10 (Fig. 1a). This revealed a potential clash between hFN10 and αIIb propeller involving Ser^1500^-Lys in the C-terminal F-G loop of hFN10 and Val^156^-Glu in the longer helix-containing D2-A3 loop of αIIb. In addition, the hFN10 ligand Arg^1493^ could not make the critical bidentate salt bridge with αIIb-Asp^224^. We therefore substituted Ser^1500^-Lys in hFN10 with Gly, and replaced the ligand Arg^1493^ with the longer L-homoarginine (Har), changes that we predicted would not adversely affect FN10 folding or the π-π interaction between β3-Tyr^122^ and Trp^1496^ of the modified peptide Hr10. The presence of Har in Hr10 was confirmed by Mass spectroscopy (Supplementary Figure 1).

**Figure 1.**
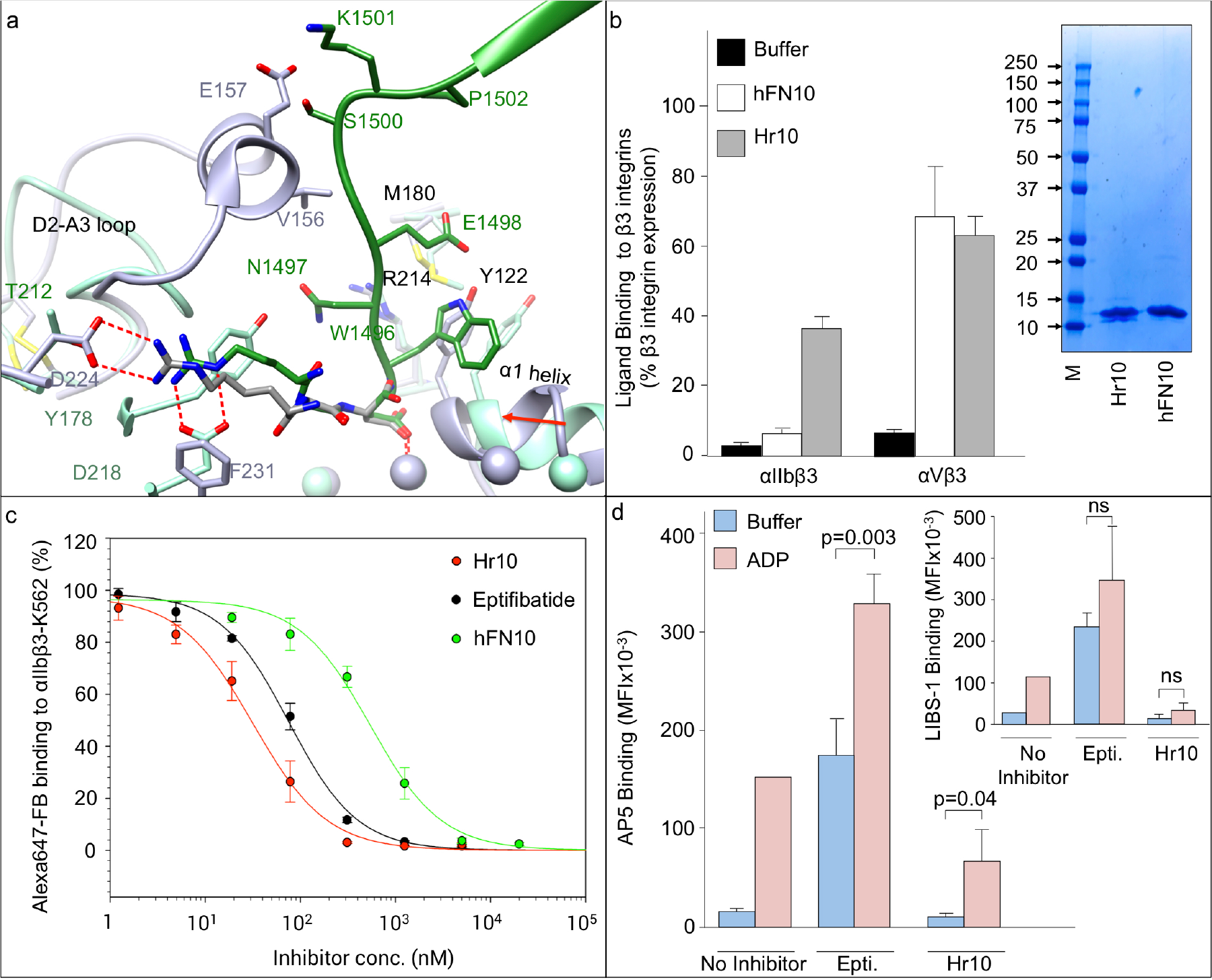
Structure-guided design and binding properties of Hr10. a) Ribbon diagrams of hFN10/αVβ3 (light green) and Eptifibatide/αIIbβ3 (light purple) crystal structures superposed on the βA domain of each, with the metal ions at LIMBS, MIDAS and ADMIDAS shown as spheres in the respective colors. Relevant segments of the propeller and βA domains and of hFN10 (dark green) and Eptifibatide (dark gray) are shown. The MIDAS ion is ligated by the aspartate residue of each ligand. Residues (single letter code) specific to each structure are shown in the respective color, with residues or loops common in both shown in black. Oxygen, nitrogen, and sulfur atoms are in red, blue and yellow, respectively. The inward movement (red arrow) of the α1 helix (and ADMIDAS ion) towards MIDAS, driven by binding of the partial agonist Eptifibatide to αIIbβ3, is absent in hFN10-bound αVβ3, the result of a π-π interaction between ligand W^1496^ and β3-Y^122^. β3-R^214^ and β3-M^180^ contribute to the stability of ligand W^1496^. The homoarginine from Eptifibatide forms a bidentate salt bridge with αIIb-D^224^, whereas R^1493^ of hFN10 contacts αV-D^218^ (replaced by F^231^ in αIIb). A clash between the c-terminal F-G loop of hFN10 and the longer D2-A3 loop of αIIb propeller, replacement of D^218^ in αV with F^231^ in αIIb and the shorter side chain of R^1493^ (vs. homoarginine in Eptifibatide) are predicted to account for the poor binding of hFN10 to αIIbβ3. b) FACS analysis showing binding of Hr10 and hFN10 (each at 1.5 μM) to αIIbβ3-K562 (αIIbβ3) and αVβ3-K562 (αVβ3)(mean±S.D., n=3 independent experiments). Bound ligand was detected with a labeled anti-His tag monoclonal antibody. Binding of the anti-His alone is indicated as buffer. Inset, Coomassie stain of 10-20% SDS PAGE showing purified Hr10 and hFN10 (8 μg in each lane). MW markers (in kDa) are indicated. c) Dose response curves (mean±S.E., n=3 independent experiments) generated from FACS analyses showing displacement of Alexa-647 labeled fibrinogen (FB) bound to preactivated αIIbβ3-K562 in the presence of increasing concentrations of unlabeled Hr10, Eptifibatide or hFN10. The MFI values from the three separate FACS analyses were normalized individually before averaging as described in Methods. The IC_50_ values are stated in the text. d) Histograms (mean+S.D., n=3 experiments) showing effects of Hr10 vs. Eptifibatide (each at 1.5 μM) on integrin conformational changes. Binding of the activation-sensitive mAb AP5 or the extension-sensitive mAb LIBS-1 (inset) to human platelets in the absence or presence of ADP (5μM) was assessed following flow cytometry. Binding of the two conformation-sensitive mAbs to ADP-activated platelets in the absence of the inhibitors was often complicated by platelet aggregation, thus only single measurements could be obtained for each mAb. ns, not significant.

### Hr10 is a pure antagonist of αIIbβ3

Purified Hr10, but not hFN10, bound K562 cells stably expressing recombinant αIIbβ3 (αIIbβ3-K562) (Fig.1b) and maintained its binding to αVβ3-K562 (Fig.1b), as expected. Hr10 inhibited binding of Alexa647-labeled soluble FB to activated αIIbβ3 more effectively than Eptifibatide (IC_50_ 30.3±4.8 nM [mean ± S.E., n=3] vs. 73.2 ± 7.0 nM for Eptifibatide, p=1.79×10^−5^) (Fig.1c). hFN10 bound minimally to activated αIIbβ3-K562, with an order of magnitude higher IC_50_ of 474.0±73.4 nM (Fig. 1c).

Binding of Eptifibatide (1.5 μM) to human platelets in the absence or presence of 5 μM ADP induced conformational changes in αIIbβ3 reported by binding of the activation-sensitive and extension-sensitive mAbs AP5 and LIBS-1, respectively ^24^ (Fig. 1d). In contrast, binding of Hr10 (1.5 μM) did not induce these changes, and suppressed AP5 and LIBS-1 binding to ADP-activated platelets (Fig. 1d). Thus Hr10 acts as a pure antagonist of αIIbβ3.

### Crystal structure of Hr10/β3 integrin complex

To elucidate the structural basis of pure antagonism, we determined the crystal structure of the Hr10/integrin complex at 3.1Å resolution (Fig. 2a and Supplementary Table 1) by soaking Hr10 into preformed αVβ3 ectodomain crystals (crystal packing of the αIIbβ3 ectodomain does not allow access of large ligands to MIDAS). The structure confirmed presence of the homoarginine at position 1493 in Hr10. Har^1493^ forms a bidentate salt bridge with αV-Asp^218^ and a cation-π interaction with αV-Tyr^178^, but does not contact αV-Thr^212^ (which replaces αIIb-Asp^224^). The ligand Asp^1495^ directly coordinates the metal ion at MIDAS, with Trp^1496^ making a π-π interaction with βA-Tyr^122^, stabilized by an S-π interaction with βA-Met^180^ (Fig. 2a) and a critical hydrogen bond between the carbonyl of Trp^1496^ and Nε of βA-Arg^214^. Bound Hr10 prevented the activating inward movement of the α1 helix (reported by β3-Tyr^122^) towards MIDAS, and the conformational changes at the C-terminal end of βA domain, which trigger integrin extension. Superposition of the βA domains from the Hr10/αVβ3 and Eptifibatide/αIIbβ3 structures (Fig. 2b) shows that the Ser^1500^-Lys/Gly substitution removes the predicted clash with the αIIb propeller. The Nε, Nh1 and Nh2 amino groups of the ligand Har^1493^ superpose well on those of Har^2^ in Eptifibatide, and could likewise form the critical bidentate salt bridge with αIIb-Asp^224^, accounting for the high affinity binding of Hr10 to αIIbβ3. β3-Tyr^122^ is replaced with Phe^122^ in mouse β3, and the stabilizing salt bridge β3-Arg^214^ makes with β3-Asp^179^ is replaced with a H-bond with β3-Asn^179^ in mouse, both substitutions likely contributing to the poor binding of Hr10 to mouse αIIbβ3 (not shown).

**Figure 2.**
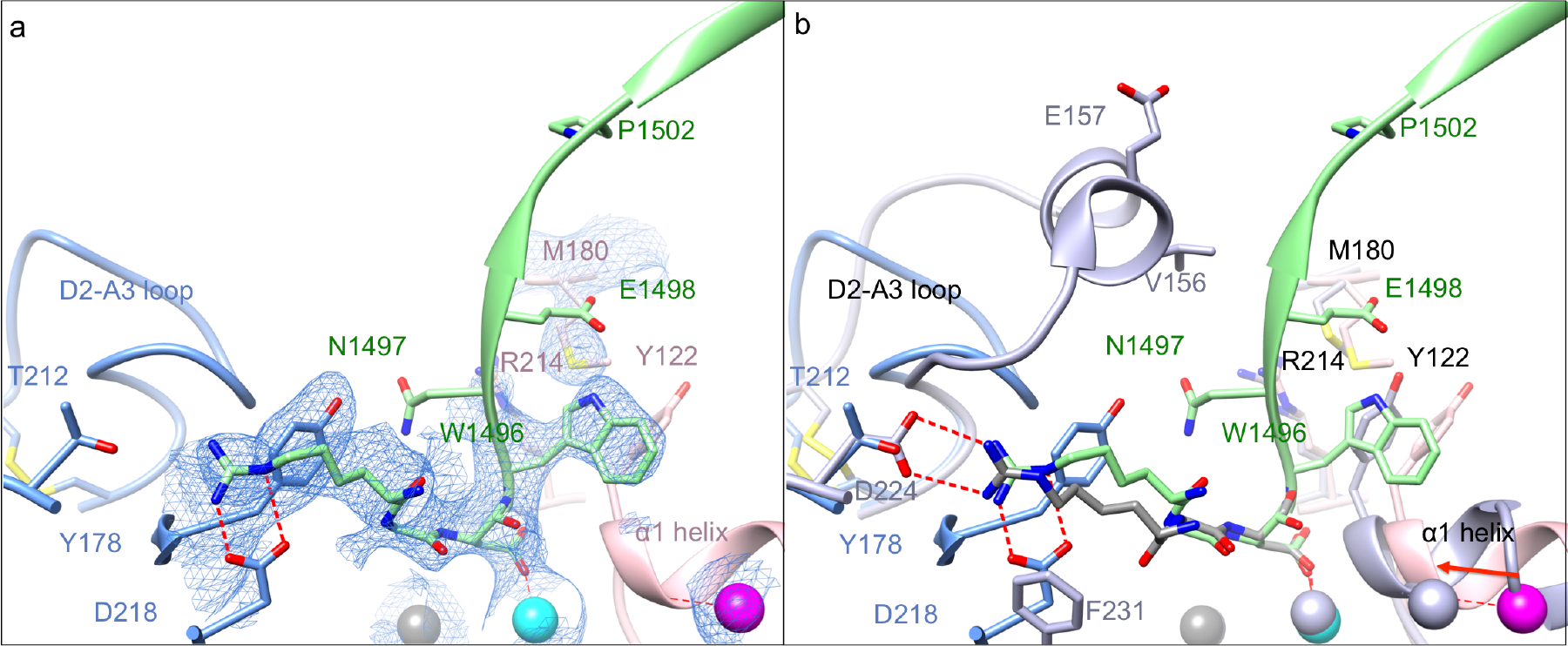
Crystal structure of Hr10/β3 integrin complex. a) Ribbon diagram of the crystal structure of Hr10/αVβ3 complex (same view as Figure 1a) showing the electron density map at 1.0 σ (blue mesh) of the ligand-binding region. Relevant portions of Hr10 (light green), αV propeller (light blue) and the β3A domain (rose color) are shown. Side-chains are shown as sticks in the respective colors. The Mn^2+^ ions at LIMBS, MIDAS and ADMIDAS are in grey, cyan, and magenta spheres, respectively. Oxygen, nitrogen, and sulfur atoms are colored as in Figure 1a. Water molecules are not shown. Hr10’s W^1496^ forms a π-π interaction with β3-Y^122^, and Har^1493^ forms a bidentate salt bridge with αV-D^218^. b) Ribbon diagrams of the crystal structures of Hr10/αVβ3 (light green) and Eptifibatide/αIIbβ3 (light purple) superposed on the βA domain of each. View, domain, side chain and metal ion colors are as in 2a. Note the removal of the predicted clash of Hr10 with D2-A3 loop of αIIb and predicted formation of Har^1493^-αIIb-D^224^ salt bridge.

### Effects of Hr10 on human platelet aggregation and secretion

Hr10 blocked platelet aggregation induced by the agonists collagen, ADP and TRAP as effectively as Eptifibatide (Fig. 3a-d). The adenine nucleotides ADP and ATP are co-released from dense (δ-) granules during platelet activation, and interact with the platelet P_2_ receptors to amplify ongoing platelet activation. Both Hr10 and Eptifibatide (at 1.5 μM) inhibited ADP (20 μM)-induced ATP secretion from dense (δ-) granules in whole blood by 71 ± 14 % and 60 ± 20 %, respectively (Fig. 3e), but did not significantly alter ADP-induced secretion from human platelet α-granules (reported by CD62P expression) or from lysosomes (reported by CD63 expression) (Fig. 3f), as noted earlier for Abciximab ^26^ and Tirofiban ^27^.

**Fig. 3.**
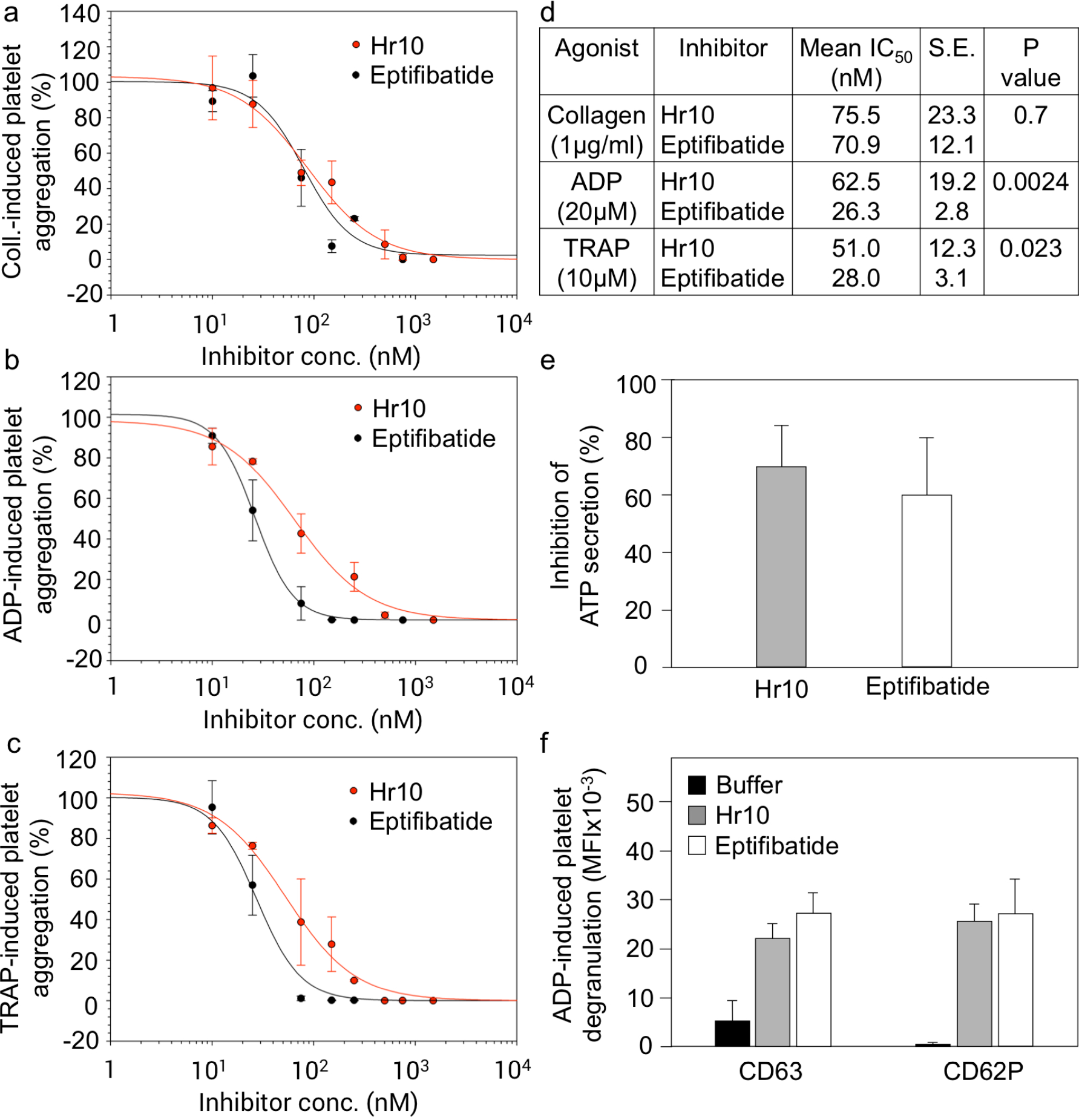
Effect of Hr10 and Eptifibatide on platelet aggregation and secretion. (a-c) Dose response curves (mean ±S.E., n=3 experiments from 3 different donors) showing effects of the inhibitors on aggregation induced by collagen (2μg/ml) (a), ADP (20 μM) (b), or TRAP (10 μM) (c). Points for the integrated impedance from the three experiments were individually normalized prior to averaging and are displayed with least-squares fits to the mean values. The respective IC_50_, S.E. and p-values from a Fisher test are listed in (d). e-f) Histograms (mean±S.D., n=3) showing the effect of Hr10 and Eptifibatide (each at 1.5 μM) on ADP (20 μM)-induced ATP secretion (e; p=0.5) and surface expression of CD63 and CD62P (f) in human platelets. No differences in expression of CD63 (p=0.15) or CD62P (p=0.72) were found in platelets exposed to Eptifibatide or Hr10.

### Hr10 preserves thrombin-induced clot retraction

Clot retraction normally helps secure hemostasis *in vivo* as evidenced by increased bleeding in mice with impaired clot retraction ^28^, or in recipients of any of the three anti-αIIbβ3 drugs ^3,29,30^. We compared the effects of Hr10 and Eptifibatide on thrombin-induced clot retraction in fresh human platelet-rich plasma (PRP) ^31^. The kinetics of clot retraction were determined from quantification of serial images of the reaction acquired every 15 minutes for the 2-hour duration of the assay. As shown in Fig.4 a, b, Hr10 did not inhibit clot retraction vs. buffer alone (p= 0.125). In contrast, Eptifibatide significantly blocked clot retraction vs. buffer (p=4.5×10^−5^), as previously shown ^29,32^. It has been reported that that αIIbβ3 antagonists that block platelet aggregation but not clot retraction exhibit affinities to inactive αIIbβ3 that are 2-3 logs lower than those to the active integrin ^2^. This was not the case with Hr10, however, as its binding to inactive αIIbβ3 (IC_50_=58.8±24.1 nM, 9 determinations from 5 experiments) was not significantly different from that to active αIIbβ3 (IC_50_ 35.2±5.7 nM, n=3 experiments; p=0.54)(Fig. 4c), and are also comparable to the affinity of Eptifibatide to αIIbβ3 on resting platelets (*k*_*D*_=120 nM)^33^.

**Figure 4.**
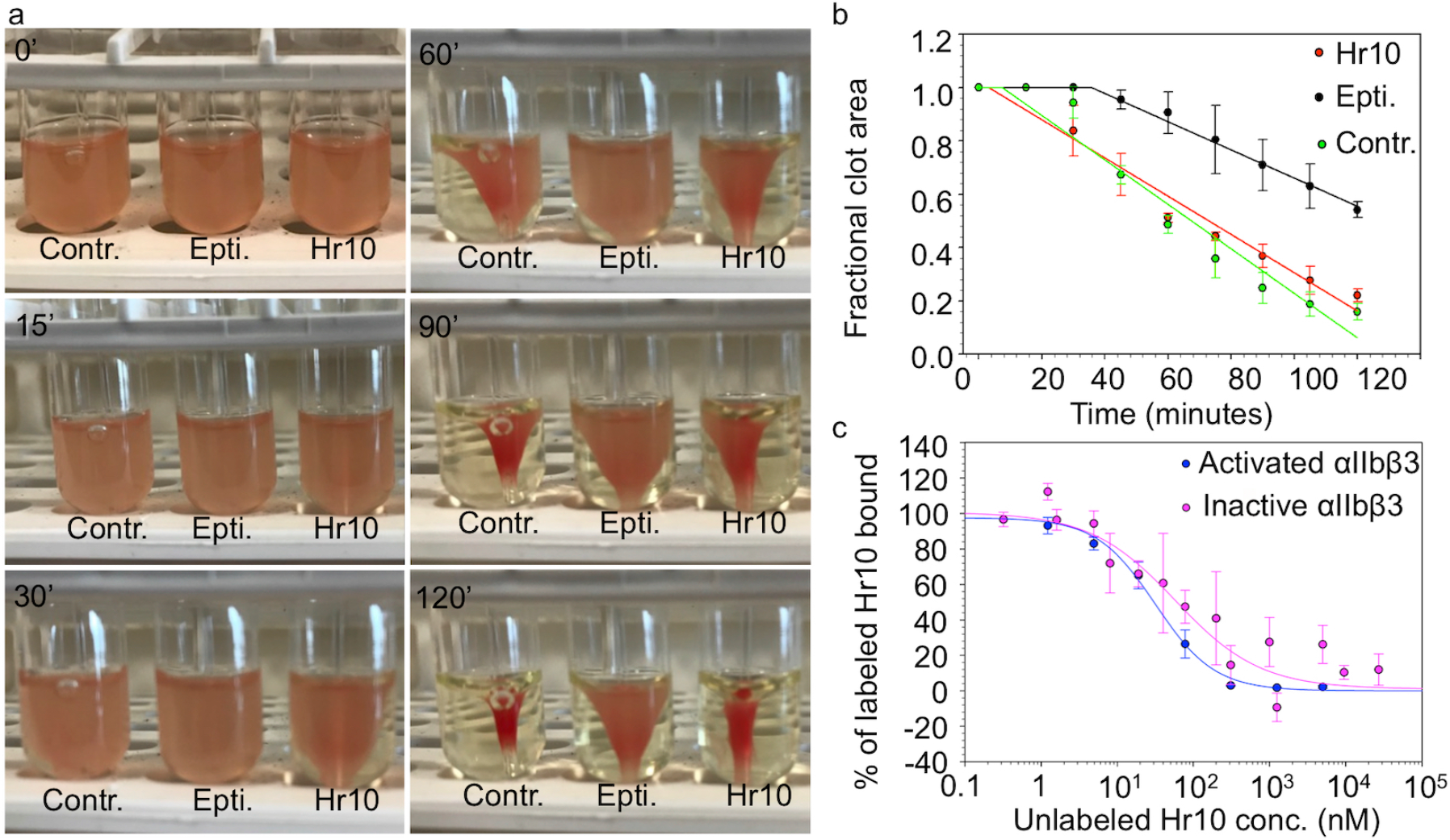
Effects of Hr10 on thrombin-induced clot retraction and binding to inactive and active αIIbβ3. a) Kinetics of clot retraction in the absence (Contr.) and presence of Hr10 or Eptifibatide (Epti.) from one representative experiment of three conducted. Clot retraction took place around a central glass rod. 5 μl of red blood cells were added per 1 ml reaction to enhance the color contrast for photography. Photographs shown were taken at 0, 15, 30, 60, 90 and 120 minutes after addition of thrombin. (B) Time course (mean ±S.E.) from three clot retraction experiments including the one shown in (a). The plot shows the fractional area occupied by the clot at 15-minute intervals with a linear regression through the points. No significant differences (p=0.125) were found in kinetics of clot retraction in buffer vs. Hr10. A lag period is noted with Eptifibatide and clot retraction was significantly reduced vs buffer (p=4.5×10^−15^). (c) Dose response curves comparing displacement of Alexa488-labeled Hr10 binding to inactive and PT-25-activated αIIbβ3 on K562 cells by increasing concentrations of unlabeled Hr10. Cell binding was analyzed by FACS. The mean fluorescence intensity values for individual Hr10 experiments (four independent experiments and 6 determinations) were initially fit with a binding curve to determine minimum and maximum MFI values to use in scaling the data. The points and error bars indicate the mean and standard error for the scaled data. The red and black lines are a least squares fit to the averages. No differences were found (p=0.54).

### Hr10 blocks microvascular thrombosis without increasing bleeding in humanized mice

To evaluate the effects of the peptides Hr10 and Eptifibatide on nascent thrombus formation under flow, we induced thrombin-mediated arteriolar injury in a humanized mouse model that predicts clinical efficacy of anti-platelet agents ^34^. NSG (NOD-*scid-IL-2Rγ*^*null*^) mice were made homozygous for human von Willebrand factor R^1326H^ (vWF^RH/RH^) ^34^, a substitution that switches binding of vWF from mouse to human glycoprotein (GP) Ib/IX, which accounts for the increased bleeding risk in these mice unless mice are infused with human platelets. To assess the effects of Hr10 and Eptifibatide on bleeding, each inhibitor was given to mice infused with human platelets (Eptifibatide ^34^, like Hr10, binds poorly to mouse αIIbβ3). Hr10 in equimolar concentrations to Eptifibatide was as effective in completely preventing nascent occlusive thrombus formation at multiple sites of laser-induced arteriolar injury (Fig. 5a). Significantly, however, and in contrast to Eptifibatide, Hr10 did not increase bleeding in the humanized vWF^*RH/RH*^NSG mice (Fig.5b).

**Figure 5.**
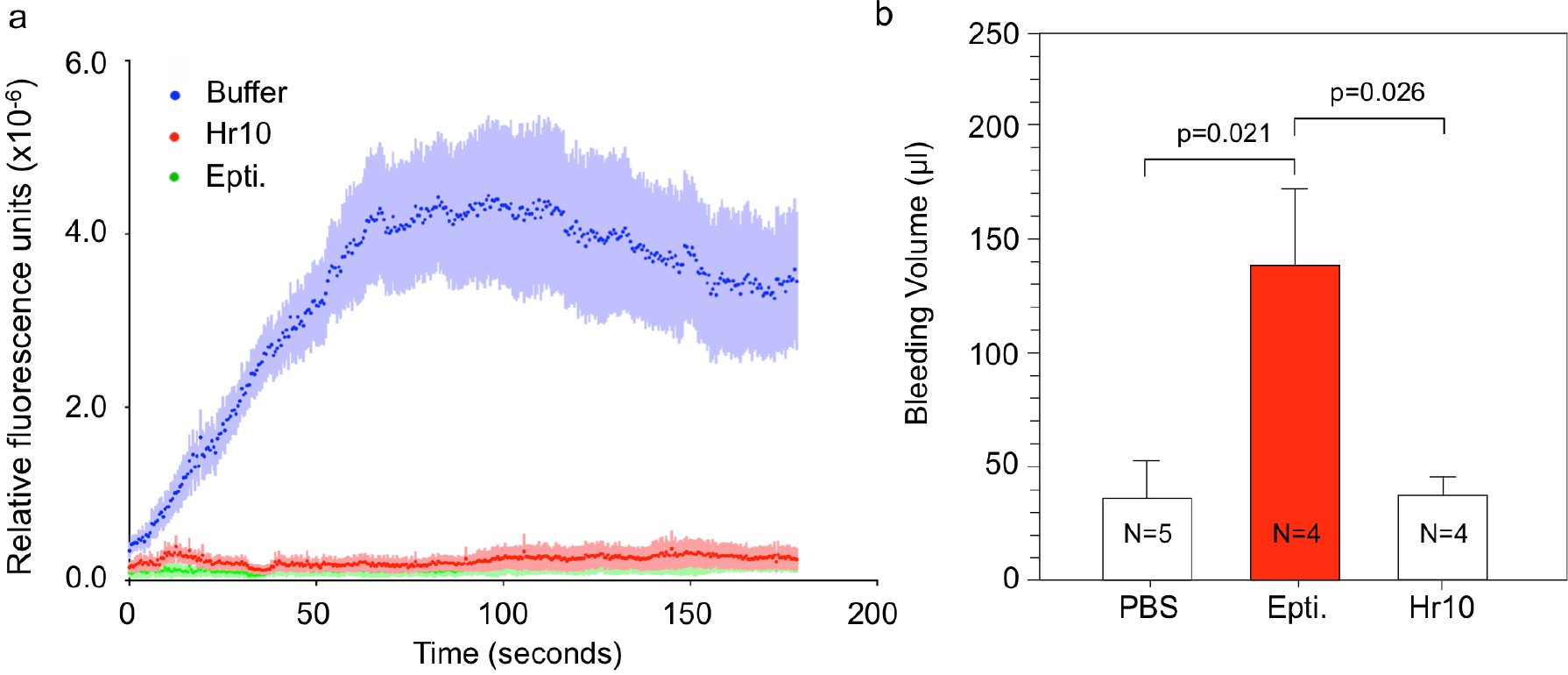
Hr10 inhibits laser-induced microvascular injury and preserves hemostasis in mice. a) Graphs showing kinetics (mean±S.E., n=4 mice with laser-induced injuries at 8 different sites made in each) of human platelet accumulation at nascent injuries during infusion of buffer (PBS) or equimolar amounts of Hr10 or Eptifibatide. *n*=4 animals per arm. There was no significant difference in human platelet accumulation in thrombi, between Hr10 and Eptifibatide-treated mice at each time point. b) Histograms (mean+S.E.) showing baseline bleeding volume in vWF^*RH/RH*^NSG mice infused with human platelets before (PBS) or after administration of Eptifibatide (Epti.) or Hr10. Epti. caused excessive bleeding (~10% of blood volume of a normal mouse). This was completely averted in presence of Hr10. p-values are indicated p=0.94 between PBS and Hr10 receiving mice). Other p-values are shown.

### Structure-guided conversion of Tirofiban into a pure αIIbβ3 antagonist

We next explored the feasibility of converting the αIIbβ3-specific partial agonist drug Tirofiban (molecular weight of 495.08) into a pure antagonist, guided by the crystal structure of Hr10/αVβ3 complex. Superposing the βA domains of Hr10/αVβ3 and Tirofiban/αIIbβ3 (2vdm.pdb) ^35^ structures show that the acidic moiety of each ligand and the following amide and sulfonamide moieties are nearly superimposable (RMSD = 0.9648)(Fig.6a), suggesting that replacing the butane-sulfonamide moiety of Tirofiban with an indole group could create the critical π-π interaction with βA-Tyr^122^. We selected the indole derivative benzoxazole in order to stabilize this interaction further by formation of a hydrogen bond between the benzoxazole oxygen and Nε of β3-Arg^214^ as in the Hr10/αVβ3 structure. A structure of such modified Tirofiban (M-Tirofiban) in complex with inactive αIIbβ3 (3fcs.pdb) was then modeled in Coot ^36^ by geometry minimization with a library generated by eLBOW in Phenix ^37^. In this model (Fig.6a), the RGD-like moiety of M-Tirofiban superimposes nicely on that of Tirofiban, with the benzoxazole moiety forming a π-π contact (4.4Å) with β3-Tyr^122^, and the benzoxazole oxygen forming a hydrogen bond (3.2Å) with Nε of β3-Arg^214^, arrangements predicted to freeze the integrin in the inactive conformation. To test this prediction experimentally, we determined the binding properties of the synthesized M-Tirofiban in cell-based assays M-Tirofiban blocked FB binding to preactivated cellular αIIbβ3 (Fig.6b) and prevented ADP-induced human platelet aggregation (Fig.6c) in the low nanomolar range (~18-30 nM), compared with 1.5-2 nM for Tirofiban. The ~10-fold reduction in affinity of M-Tirofiban vs. Tirofiban likely reflects weaker H-bonding of the benzoxazole oxygen vs. the sulfonamide oxygen of Tirofiban with Nε of β3-Arg^214^, and perhaps loss of hydrophobic contacts of the deleted butane moiety with the integrin. Nevertheless, the affinity of M-Tirofiban calculated in these assays was equivalent to that of the drug Eptifibatide (Fig.6b, c).

**Figure 6.**
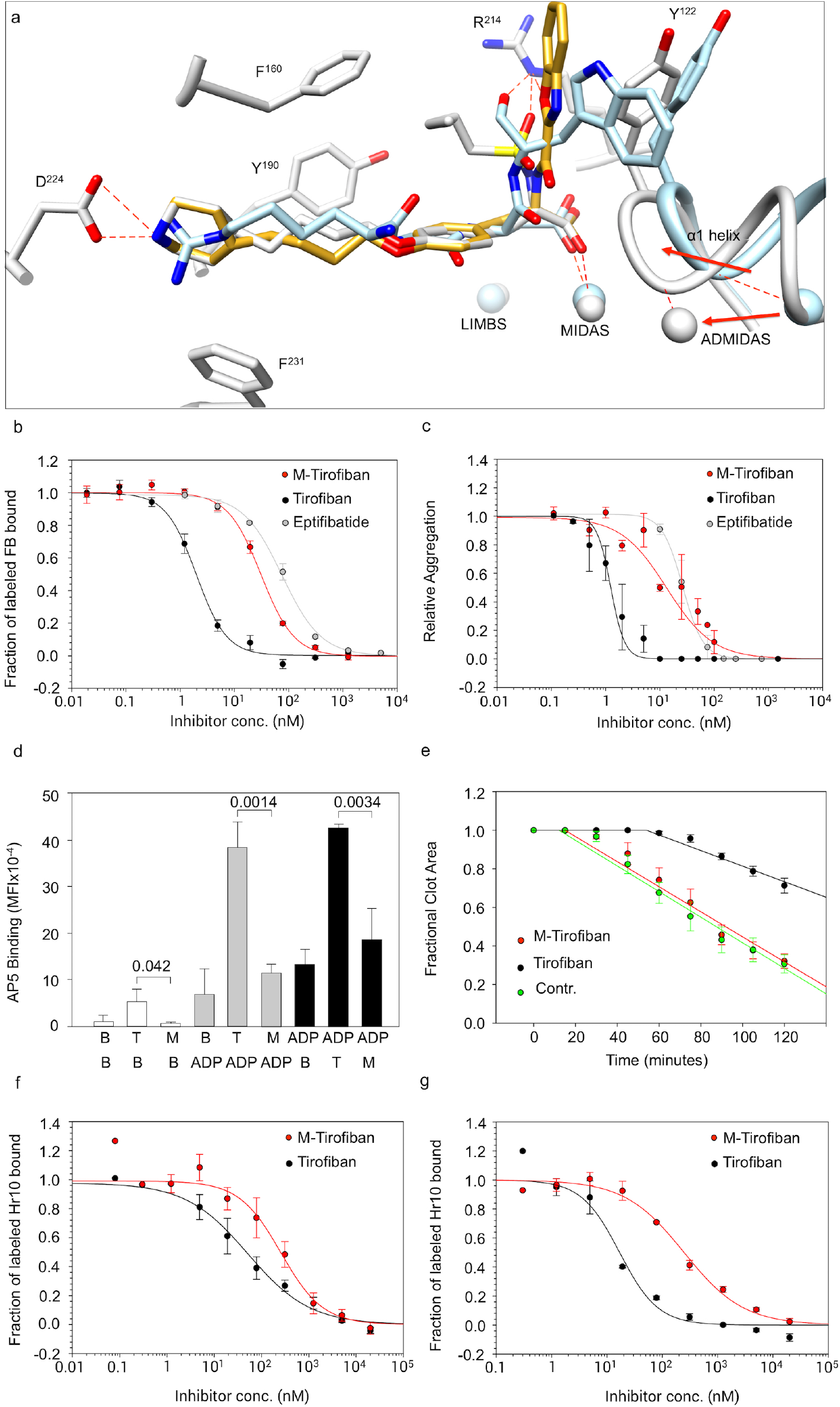
Engineering and binding properties of M-Tirofiban. a) A model of bound M-Tirofiban (gold) superposed on the crystal structures of Tirofiban/αIIbβ3 (gray) and Hr10/αVβ3 (light blue). The βA domain of each was used in superposition. The metal ions at LIMBS, MIDAS and ADMIDAS and relevant residues are shown in the respective colors. Contacts are shown as dotted red lines. See text for details. b) Dose response curves (mean±S.E., n=4 experiments) showing displacement of labeled fibrinogen bound to preactivated αIIbβ3-K562 by Tirofiban or M-Tirofiban yielding IC_50_s of 1.98±0.19 nM, and 30.9±3.3 nM, respectively. Displacement of labeled fibrinogen by Eptifibatide (Fig. 1c) is added for comparison. c) Dose response curves (mean ±S.E., n=5 experiments) showing effects of Tirofiban or M-Tirofiban on human platelet aggregation from three different donors induced by ADP (20 μM) yielding IC_50_s of 1.41±0.23nM and 18.5±5.4nM, respectively. Effect of Eptifibatide on platelet aggregation (Fig. 3b) is shown for comparison. d) Histograms (mean±S.D., n=3 independent experiments) showing binding of AP5 mAb to human platelets in presence of buffer (B), Tirofiban (T)(150 nM) and M-Tirofiban (M)(1.5 μM) (white histograms) alone, and before (gray histograms) or after (black histograms) addition of ADP (5μM). Numbers represent p-values. No significant differences were found between B and M before (p=0.273) or after (p=0.81) ADP addition. e) Kinetics of clot retraction in the absence or presence of Tirofiban, or M-Tirofiban (mean ±S.E.) from three experiments. Kinetics of clot retraction was not different between buffer control (contr.) and M-Tirofiban (p= 0.61). (f, g) Dose response curves (mean+S.E.) comparing displacement of Alexa488-labeled Hr10 binding to inactive (f, n=5) and mAb PT-25-activated αIIbβ3-K562 (g, n=3) by increasing concentrations of Tirofiban or M-Tirofiban in presence of physiologic concentrations of Mg^2+^ and Ca^2+^ (1 mM each). The respective IC_50_s were 51.3±19.2nM, and 257.2±88.0 nM for inactive and 16.9±2.4 nM and 247.1±29.3 nM for active αIIbβ3. The lower affinities of both compounds are explained by the requirement for more inhibitor to displace high affinity binding of Hr10 (compared to fibrinogen in b) to αIIbβ3.

Binding of Tirofiban to αIIbβ3 at the clinically effective concentration of 150 nM ^19^ induced expression of the AP5 epitope, which was markedly increased upon addition of ADP or when Tirofiban is added after platelet exposure to ADP (Fig.6d). In contrast, M-Tirofiban (at the equipotent concentration of 1.5μM) did not induce AP5 expression, prevented that induced by subsequent addition of ADP and even suppressed ADP-induced AP5 expression when M-Tirofiban was added afterwards (Fig.6d). And while Tirofiban effectively blocked thrombin-induced clot retraction, M-Tirofiban did not (Fig.6e) and showed equivalent binding affinities to inactive and active αIIbβ3 (Fig.6f, g).

## Discussion

The Achilles heel of current anti-thrombosis drugs that directly target αIIbβ3 is serious bleeding, an adverse outcome that remains high with use of the newer inhibitors of P_2_Y_12_ and thrombin receptors ^38,39^. Several attempts are therefore being made to develop new anti-thrombosis drugs that maintain efficacy but preserve hemostasis. These include targeting collagen receptors α2β1 and GPVI ^40,41^, accelerating ADP degradation with CD39 ^42^ or interfering with ADP-induced cell signaling with a PI3Kβ inhibitor ^43^. However, these approaches do not affect platelet activation induced by other potent agonists, and some targets (e.g. PI3Kβ and α2β1) are not platelet-specific. Platelet-leukocyte interactions are also being targeted: interfering with binding of leukocyte integrin CD11b to platelet GP1bα delayed thrombosis without prolonging bleeding time in normal mice ^44^. However, platelet-leukocyte interactions are mediated by multiple receptor-counterreceptor pairs, the relative importance of which may vary with the nature of the pathologic state. Two recent attempts targeted αIIbβ3 more directly. In one approach, a short cytoplasmic β3-derived peptide inhibited αIIbβ3 outside-in signaling and prevented thrombosis without prolonging bleeding time, but is not β3-integrin specific ^32^. The second approach utilized low affinity non-RGD small molecules RUC2 and RUC4 that engage the arginine pocket in αIIb ^45^ and prevent FeCl3-induced thrombotic arterial occlusion in mice but its effects on clot retraction or bleeding were not reported ^46^.

The present data show that pure orthosteric antagonists of αIIbβ3 block arteriolar thrombosis while preserving hemostasis, thus demonstrating that partial agonism and antagonism are not inseparable. Pure orthosteric antagonism of αIIbβ3 offers significant advantages over the other approaches aimed at preserving hemostasis. First, by targeting αIIbβ3 MIDAS directly, pure antagonists block binding of several prothrombotic ligands, some of which like CD40L ^47^ bind leukocyte CD11b ^48^ and thus contribute to platelet-leukocyte interactions. Second, these high affinity pure orthosteric inhibitors do not induce the conformational changes directly and block these when induced by inside-out integrin activation. Third, Hr10, a minor variant of a human natural ligand, is expected to be minimally immunogenic. Whether preformed antibodies that recognize αIIbβ3 in complex with Tirofiban ^19^ no longer do so when αIIbβ3 is bound by M-Tirofiban or Hr10 can now be tested. Success in converting the partial agonist Tirofiban into a pure antagonist using the Hr10/integrin structure also underscores the primacy of the stable π-π Trp^1496^ - β3-Tyr^122^ contact in preventing the activating global conformational change in αIIbβ3, and suggest that this approach may be applicable to engineering drug candidates targeting other integrins, where inadvertent conformational changes may also compromise patient safety.

The precise mechanism by which prevention of the agonist-induced conformational changes in αIIbβ3 by pure antagonists results in preservation of clot retraction is presently unknown, but there are several possibilities. Clot retraction occurs in response to the binding of polymeric fibrin to αIIbβ3, thus linking the integrin to actomyosin ^49^. When compared with fibrinogen, polymeric fibrin binds αIIbβ3 with higher affinity ^50^. So one possibility is that high affinity of the partial agonists is necessary to block fibrin-αIIbβ3 interaction and hence clot retraction. This is unlikely since Hr10 and Eptifibatide have comparable affinities in blocking soluble fibrinogen binding to activated αIIbβ3 and agonist-induced platelet aggregation. Preservation of clot retraction by the pure antagonists could not be explained either by a weaker affinity to inactive αIIbβ3 ^51^, since affinities of the pure antagonists to inactive and active αIIbβ3 are similar. A recent study showed that fibrin binds αIIbβ3 even when all the RGD motifs in fibrin are deleted ^50^, reflecting presence of MIDAS-independent fibrin binding sites, localized recently to the αIIb propeller domain ^52^. Since αIIbβ3 on non-activated platelets binds surface-immobilized fibrin ^53,54^, it is expected to also do so when occupied by Hr10 (or M-Tirofiban). It has been shown that αIIbβ3-dependent fibrin clot retraction kinetics correlates with intracellular protein tyrosine dephosphorylation, which is inhibited by binding of Eptifibatide or Abciximab to αIIbβ3 ^29^. The availability of the pure orthosteric inhibitors of αIIbβ3 should now provide a vital tool to dissect the mechanisms linking integrin conformation to clot retraction.

The dual specificity of Hr10 to both β3 integrins is shared with the drug Abciximab ^55^, a property thought to contribute to the long-term clinical benefits of Abciximab in acute coronary syndromes ^56,57^. In addition, dual specificity to both β3 integrins has shown a wide range of anticancer effects (reviewed in ^58^). For example, Abciximab was effective at blocking tumor growth and angiogenesis through targeting the interaction of tumor cells with platelets and endothelial cells, in addition to direct effects on the tumor tissue ^59–62^. Hr10 may thus offer an attractive clinical candidate with minimal bleeding risk and an expected low to absent immunogenicity.

## Supporting information

Supplementary File_BioRxiv

## Acknowledgements.

We thank Drs. Donald W. Landry and Shi-Xian Deng (Columbia University, NY, NY) for synthesis of M-Tirofiban and Jennifer Cochran (Stanford University, CA) for providing K562 cells stably expressing αIIβ3 (αIIβ3-K562). We also thank Drs. Mark Ginsberg (UC San Diego, CA), Thomas J Kunicki (Scripps Research Institute, CA), Kensaku Sakamoto for providing the B-95ΔA cells and pCDF5-Har plasmid, Makoto Handa (Keio University, Tokyo, Japan) for providing mAbs LIBS-1, AP5 and PT-25-2, respectively. This work was supported by NIH grants DK088327 and DK48549 and the RICBAC Foundation (to MAA), DK101628 (to JVA) and R01HL142122 and P01HL040387 (to MP) from the National Institutes of Diabetes, Digestive and Kidney diseases (NIDDK) and Heart, Lung and Blood (NHLBI) of the National Institutes of Health and grants from National Core Facility for Biopharmaceuticals, Ministry of Science and Technology, Taiwan (S-W Lin and I-Su).

## Author contributions

MAA conceived and oversaw all experiments. BLD performed the aggregation and clot retraction assays. JLA designed and performed the ligand binding studies. JVA collected the diffraction data and refined the structures. JVA, JXP and MAA performed model building and structure analysis. S-W Lin and I-Su generated the vWF^RH/+^NSG mice. VH, HSA and MP performed the mouse studies. All authors interpreted data. MAA had the primary responsibility for writing the manuscript.

## Methods

### Reagents and antibodies

Restriction and modification enzymes were obtained from New England Biolabs Inc. (Beverly, MA). Cell culture reagents were purchased from Invitrogen (San Diego, CA) or Fisher Scientific (Hampton, NH). The Fab fragment of AP5 was prepared by papain digestion followed by anion exchange and size-exclusion chromatography. Hybridoma producing β3 conformation-insensitive mAb AP3 was bought from ATCC and antibody purified by affinity chromatography. Alexa Fluor 488-conjugated mAbs against human CD62P and CD63 were bought from Santa Cruz Biotechnology, Dallas, TX. Alexa Fluor647-conjugated anti-human CD42b mAb was from R&D Systems, Minneapolis, MN. APC-labeled goat anti-mouse Fc-specific antibody was from Jackson ImmunoResearch (West Grove, PA). Alexa Fluor 647-conjugated Penta-His mAb was purchased from Qiagen, Germantown, MD. Eptifibatide and Tirofiban were purchased from Millipore-Sigma (Burlington, MA). M-Tirofiban [OC(=O)[C@H](CC1=CC=C(OCCCCC2CCNCC2)C=C1)NC(=O)C1=NC2=C(O1)C=C C=C2] was synthesized at the Organic Chemistry Collaborative Center, Columbia University Irving Medical Center, NY. Purity for M-Tirofiban was determined through high-performance liquid chromatography and found to be > 95% pure. The plasmid pCDF5-Har, containing two copies of a UAG recognizing tRNA and the tRNA synthase (Har-Rs) for charging UAG tRNAs with Har, and the *E. coli* strain B-95ΔA containing a deletion of release factor 1 (*prfA)* and 95 synonymous TAG stop codon mutations, were kindly provided Dr. Kensaku Sakamoto (RIKEN, Yokohama, Japan) ^63^. L-Har and TRAP-6 were purchased from Bachem. ADP, collagen, ATP, Chrono-luminescence reagent and human thrombin were purchased from Chrono-log (Havertown, PA).

### Plasmids, mutagenesis, protein expression, purification and mass spectrometry

Human αVβ3 ectodomain was expressed and purified as described ^24^. hFN10 was expressed in BL21-DE3 bacteria and purified by affinity chromatography followed by gel filtration as described ^24^. hFN10 containing a TAG stop codon at position 1493 was generated by PCR-based mutagenesis with the Quick-change kit (Agilent Technologies), cloned into bacterial expression plasmid pET11a and verified by DNA sequencing. A bacterial stock of *E. coli* strain B-95ΔA containing plasmids pCDF5-Har and pET-11a/hFN10-TAG grown in LB media supplemented with 5 mM L-Har, 50 μg/ml kanamycin (pCDF5-Har) and 100 μg/ml ampicillin (pET-11a) was prepared and used to express Hr10. Bacterial cultures at ~0.5 A (600 nm) were induced with 0.3 mM IPTG and grown for 8 hours at room temperature. Hr10 was purified as for hFN10 ^24^ and purity assessed by fractionation on gradient SDS-PAGE gels followed by Coomassie staining.

### Cell lines, cell culture and transfection

Human αVβ3-K562 cells have been previously described ^24^. K562 cells stably expressing αIIbβ3 (αIIbβ3-K562) were kindly provided by Jennifer Cochran (Stanford University, CA) ^64^. K562 cells were maintained in Iscove’s modified Dulbecco’s medium plus G418 (0.5–1.0 mg/ml), supplemented with 10% fetal calf serum, 2mM L-glutamine, penicillin and streptomycin.

### Ligand binding and flow cytometry

For ligand binding assays, αIIβ3-K562 or αVβ3-K562 cells (1×10^6^) were suspended in 100μl of WB (20 mM Hepes, 150 mM NaCl, pH 7.4, containing 0.1% [w/v] bovine serum albumin and 1mM each of MgCl2 and CalCl2) and incubated first with Hr10 or hFN10 (each at 3-10 μg/ml) for 30 min at room temperature (RT). After washing, cells were incubated for 30 additional minutes at 4°C with Alexa Fluor 647 conjugated Penta-His mAb. Integrin expression was independently analyzed for each condition by incubating cells with the Alexa647-conjugated AP3 (10 μg/ml) for 30 min on ice. Cells were washed, re-suspended, fixed in 2% paraformaldehyde and analyzed using FACSCalibur or BD-LSRII flow cytometers (BD Biosciences). Ligand binding was expressed as mean fluorescence intensity (MFI), as determined using FlowJo software. Mean and SD from independent experiments were calculated, and compared using Student’s t-test.

For ligand binding competition studies, 100μl of PT-25-activated αIIbβ3-K562 (1×10^6^) were incubated for 30 minutes at RT with serially diluted concentrations of Hr10, hFN10 or Eptifibatide in the presence of 0.5 μM Alexa647-conjugated FB. Cells were washed, fixed in 2% paraformaldehyde and analyzed by flow cytometry. Ligand binding was expressed as IC_50_ of cells in the absence of competitor ligands. Mean and SD from three independent experiments were calculated, and compared using Student’s t-test.

### Platelet aggregation and ATP secretion

Platelet aggregation and ATP secretion in whole blood were performed in a Chrono-Log model 700 two-channel lumi-aggregation system following the manufacturer's instructions. Blood was drawn directly into 3.2% sodium citrate from healthy volunteers after signing an informed consent form approved by the Human Subjects Committee at the Massachusetts General Hospital, and blood was used within 3 hours. None of the subjects were taking any medications for at least 10 days prior. For impedance aggregation measurements, 0.5 ml of blood was mixed with 0.5 ml physiologic saline supplemented with inhibitors and incubated at 37° C for 5 minutes without stirring. Measurements were performed with stirring at 1,200 rpm at 37° C. Values for each data point represent impedance measurements following application of agonist integrated over 5 minutes. Data points for an individual dose curve were serially collected from a single draw and analyzed with SigmaPlot (Systat Software, San Jose, CA) using a least-square fit to a logistic curve and the IC_50_ value determined from the fitted parameter. ATP secretion proceeded similarly except that 0.45 ml of whole blood were added to 0.45 ml of saline supplemented with various concentrations of Hr10 or Eptifibatide to produce the desired concentration at 1.0 ml. Following incubation for 5 minutes at 37° C, 100 μl of Chrono-lume reagent was added and aggregation initiated. The luminescence signal was quantified with a non-aggregated sample supplemented with an ATP standard.

### Binding of mAbs

αVβ3-K562 cells or transiently transfected HEK293T (0.5×10^6^ in 100 μl WB) were incubated in the absence or presence of unlabeled Hr10 or Eptifibatide, each at 1.5μM, for 20 min at RT. Alexa647-labeled AP5 Fab or unlabeled anti-LIBS-1 (each to 10 μg/ml) were added, and cells incubated for an additional 30 min before washing. Alexa647-labeled AP3 was used for normalization of integrin expression. APC-labeled goat anti-mouse Fc-specific antibody was added to anti-LIBS-1-bound cells for an additional 30 min at 4°C, cells washed and processed for flow cytometry. Binding of anti-CD62 and anti-CD63 mAbs to platelets was performed by incubating (20 min, RT) 100 μl of ligand-pretreated 3.2% sodium citrate whole blood with either Alexa488-labeled mAb (at 10 μg/ml) in the presence of 10 μg/ml Alexa647-labeled anti-CD42b. Cells were fixed in 2% paraformaldehyde and CD62 and CD63 expression was analyzed by flow cytometry in the CD42b positive population.

### Crystallography, structure determination and refinement

Human αVβ3 ectodomain was purified and crystallized by the hanging drop method as previously described ^20^. Hr10 was soaked for 3 weeks into the preformed αVβ3 crystals at 1.5 mM in the crystallization well solution containing 1 mM Mn^2+^. Crystals were harvested in 12% PEG 3500 (polyethylene glycol, molecular weight 3500), in 100 mM sodium acetate, pH 4.5, 800 mM NaCl plus 2 mM Mn^2+^ and FN10 (at 1.5 mM), cryoprotected by addition of glycerol in 2% increments up to 24% final concentration, and then flash-frozen in liquid nitrogen. Diffraction data was collected at ID-19 of APS, indexed, integrated, scaled by HKL2000 ^65^, and solved by molecular replacement using 3IJE as the search model in PHASER. The structure was refined with Phenix using translation-liberation-screw, automatic optimization of X-ray and stereochemistry, and Ramachandran restriction in the final cycle. Data collection and refinement statistics are shown in Table 1. The coordinates and structure factors of αVβ3/Hr10 have been deposited in the Protein Data Bank under accession code 6NAJ. Structural illustrations were prepared with Chimera.

### Generation of vWF^R1326H^ knock-in (KI) NSG mice

CRISPR/Cas 9 technology was used to generate the vWF R1326H KI mice of NSG background with a mutation of specific nucleotide at the exon 28 of the mouse vWF gene, resulting in replacing the Arginine (codon CGT) at amino acid no.1326 by Histidine (codon CAT). An sgRNA was designed according to the online resources, the sgRNA Designer: CRISPRko and the Cas-OFFinder, and the sgRNAs with less than 3 mismatches and less than 25 off-target sites were used. The sgRNA target sequence was 5’- CTTGAGCTCAA GGTAGGCAC -3’. The histidine codon was repaired into the gene with a single-stranded oligos. (5’- ACATCTCTCAGAAGCGCATCCGCGTGGCAGTGGTAGAGTACCATGATGGATCCCATGCTTATCTTGAGCTCAAGGCCCGGAAGCGACCCTCAGAGCTTCGGCGCATCACCAGCCAGATTA-3’(Integrated DNA technologies, Inc.). Preparation of sgRNA and Cas9 RNA for pronucleus microinjection followed instruction instructor’s manual (AmpliCap-MaxTM T7 High Yield Message Maker kit). Pronuclear microinjection was performed on fertilized eggs from NSG mice. Genotyping of founder mice was performed by PCR, TA-cloning, followed by Sanger DNA sequencing. The primer sequences for PCR genotyping were 5’- TCACTGTGATG GTGTGAACC -3‘ paring with 5‘- CTGACTATCTC ATCTCTTC -3’. PCR condition was 95°C, 5 min, followed by 35 cycles of 95 °C, 30 sec, 55 °C, 30 sec, and 72 °C, 30 sec, and a final extension at 72 °C, 7 min. TA-cloning followed an instructor’s manual (T3 Cloning kit; ZGene Biotech Inc.). Production of the vWF R1326H KI NSG mice was carried out by the Transgenic Mouse Model Core Facility.

### Clot retraction

750 μl of Tyrode’s buffer supplemented with inhibitor was mixed in a glass culture tube with 200 μl of PRP and 5 μl red blood cells. Clotting was initiated by addition of 50 μl thrombin at 10 units/ml in saline and a sealed Pasteur pipette secured in the tube center. Digital photographs of the experiment were taken at 15-minute intervals over 2 hours. Images were analyzed with ImageJ software to determine the area occupied by the clot and plasma. Plots of the relative areas and linear regressions were performed with SigmaPlot (Systat Software, San Jose, CA).

### Cremaster laser injury animal studies

Human blood was collected in 0.129 M sodium citrate (10:1 vol/vol). Blood was obtained from healthy donors after informed consent under a protocol approved by the Children’s Hospital of Philadelphia Internal Review Board in accord with the Helsinki Principles. Platelet-rich plasma (PRP) was separated after centrifugation (200*g*, 10 minutes) at room temperature (RT). The platelets were then isolated from PRP, and prostaglandin E1 (Sigma-Aldrich) added to a final concentration of 1 μM. Platelets were then pelleted by centrifugation (800*g*, 10 minutes) at RT. The pellet was washed in calcium-free Tyrode’s buffer (134 mM NaCl, 3 mM KCl, 0.3 mM NaH2PO4, 2 mM MgCl2, 5 mM HEPES, 5 mM glucose, 0.1% NaHCO3, and 1 mM EGTA, pH 6.5), and re-suspended in CATCH buffer (PBS and 1.5% bovine serum albumin, 1 mM adenosine, 2 mM theophylline, 0.38% sodium citrate, all from Sigma-Aldrich). Platelet counts were determined using a HemaVet counter (Drew Scientific). Intravital microscopy was performed as previously described ^66 67^. All animal experiments were approved by the IACUC of the CHOP, and all investigators adhered to NIH guidelines for the care and use of laboratory animals. vWF^*RH/RH*^NSG male mice were studied after being anesthetized using sodium pentobarbital (80 mg/kg) injected intraperitoneally. Mice were maintained under anesthesia with the same anesthetic delivered via a catheterized jugular vein at 5 mg/ml throughout the experiment. The cremaster muscle was surgically exteriorized and continuously superfused with a physiological buffer (PBS containing 0.9 mM CaCl_2_ and 0.49 mM MgCl_2_) maintained at 37°C throughout the entire experiment and equilibrated with a mixture of 95% N2 and 5% CO2. Human platelets, 400 million per mouse, were labeled with mouse anti-human CD41 F(ab′)2 (BD Biosciences) conjugated to Alexa Fluor-488 and infused into the jugular vein, followed by Alexa Fluor-647 rat anti-mouse CD41 F(ab′)2 (Thermo Fisher) to detect endogenous mouse platelets ^68^. Vascular injury was induced with an SRS NL100 pulsed nitrogen dye laser (440 nm) focused on the vessel wall through the microscope objective. Each injury was followed for three minutes. Eptifibatide was used at 5μg/mouse (equivalent to the clinically effective dose of ~1.5 μM ^69^) and Hr10 at the equimolar concentration (60 μg/mouse). Drugs were infused 5 minutes prior to injury via the jugular vein. Pre and post drug measurements were made in the same animal. Wide-field images of thrombi were recorded using a Hamamatsu ORCA Flash 4.0 V3 CMOS camera (Hamamatsu, Japan) coupled to an Excelitas X-Cite XLED light source. The microscope, cameras, and light sources were all controlled using Slidebook 6.0 software (Intelligent Imaging Innovations). Intensity of the fluorescent signal was used to measure incorporated platelets ^67^. Eight injuries were made in each of four mice in each group.

### Animal bleeding studies

Pentobarbital-anesthetized vWF^*RH/RH*^NSG mice were infused retroorbitally with 8×10^8^ washed human platelets in a final volume of 200 μl (so that ~40% of the circulating platelets were human). After 5 minutes PBS, 3 μM Eptifibatide or Hr10 were administered IV. After another 5 minutes, mouse tail injury was produced by amputating an 8-mm terminal tail segment using a razor blade, which was then placed in a collection tube containing sterile water at 37°C for 10 minutes. The hemoglobin level in the water was measured by a spectrophotometer, as described ^70^ with the following modifications. Briefly, the hemolyzed whole blood/water mixture was centrifuged at 21,000xg for 5 minutes. Aliquots (20 μl) of clarified, stroma-free supernatant were diluted 10-fold in a Corning 96-well micro-plate and light absorbance measured at 575 nm (Spectramax-190 plate reader, Molecular Devices, San Jose, CA). Blood loss during the 10-minute window was measured based on a previously obtained standard curve.

### Statistical calculations

Dose-response experiments for whole blood aggregation and binding to K562 cells were conducted at least three times. Curve-fitting and statistical calculations were performed in SigmaPlot. The data points from each replicate were scaled to one another by an initial fit to a sigmoidal function to determine the minimum and maximum values. Data scaled to a maximum of 1 and a minimum of 0 were combined and fit to a sigmoidal curve to determine the IC50 value. The standard error for the IC_50_ estimate was calculated using the reduced χ^2^ method. P-values comparing IC_50_s from different inhibitors were determined using the global fit function in SigmaPlot. The two data sets were fit with all parameters separate and again where the IC_50_ value is shared between the data sets. Fisher’s F statistic was calculated from the residual sum of squares and degrees of freedom for the unshared (SS_un_, DF_un_) and shared (SS_sh_, DF_sh_) with the equation F=((SS_sh_-SS_un_)/(DF_sh_-DF_un_))/(SS_un_/DF_un_) and the p-value obtained from the F distribution. Linear regression fits to data from clot retraction experiments proceeded similarly. The Holm-Sidak test following one-way ANOVA (alpha=5.0%) was used to assess if the differences in human platelet accumulation in thrombi between Hr10 and Eptifibatide-treated mice were significant. Each time point was analyzed individually, without assuming a consistent standard deviation. For the bleeding studies, the data passed the Shapiro-Wilk normality test and hence compared using the Student’s t-test. Number of mice used for bleeding studies was based on the assumption that hemostasis is preserved in 80% of Hr10-treated mice but only 5 % of Eptifibatide-treated mice (projections supported by published reports of similar studies using eptifibatide, and the predictive clot retraction data). A significance level (p value) of 0.05 is achieved using 4 animals per group, yielding 90% statistical power ^71^.

