## Supplementary File_BioRxiv for "Structure-guided design of novel orthosteric inhibitors of integrin αIIbβ3 that prevent thrombosis but preserve hemostasis"

M. Amin Arnaout

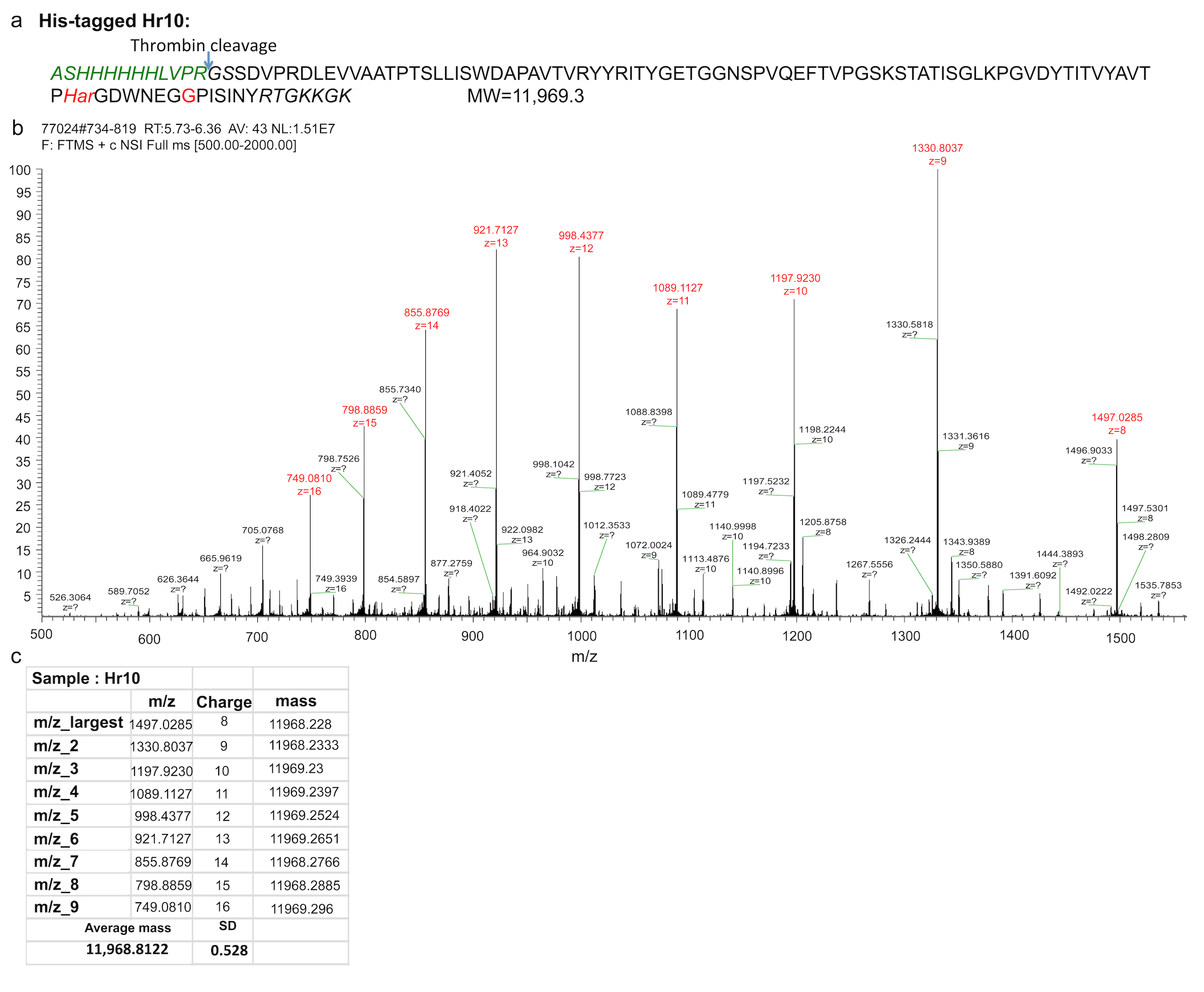
**Supplementary Figure 1**. Mass spectroscopy analysis of Hr10. a) The translated sequence of Hr10 lacking the N-terminal methionine. The homoarginine (Har) and glycine (replacing S^1500^K) residues are indicated in red. The isotopically-averaged calculated molecular weight is displayed (Protein calculator v3.4http://protcalc. sourceforge.net/cgi-bin/protcalc). b) The mass spectrum from the intact Hr10 sample. Major peaks are displayed with assigned charges. c) Table of the largest peaks showing the m/z ratios for the larges peaks, the calculated charge and resulting molecular weight. The molecular weight calculated from nine peaks is 119868.9 ± 0.5 (mean ± S.D.) as compared to the calculated weight of the protein lacking the N-terminal Met and with a single L-Har substitution (11969.3). Peaks corresponding to the protein with Arg at 1493 could not be identified (not shown).

**Supplementary Table 1. Data collection and refinement statistics.**

***Data collection*	αVβ3/Hr10**

PDB Code	6NAJ

Beamline	ID19 at APS

Space group	P3221

Unit cell dimensions (Å, ^o^)	*a*=*b*=129.7, *c*=308.2;

α=β=90, γ=120

Resolution range (Å)	50-3.1

Wavelength (Å)	0.97932

Total reflections	1,044,981

Unique reflections	55,225 (5,444)*

Completeness	100 (100)

Redundancy	8.2 (8.0)

Molecules in asymmetric unit	1

Average *I*/σ	24.9 (2.0)

*R*_merge_ (%)	9.7 (100)

*R*_meas_ (%)	10.3 (100)

*R*_sym_ (%)	3.6 (38.8)

Wilson *B*-factor	59.6

***Refinement statistics***

Resolution range (Å)	49.2-3.1

*R_factor_* (%)	24.9 (33.9)

*R_free_* (%)#	27.4 (38.9)

No. of atoms	13,498

Protein	13,137

Water	4

Mn^2+^	8

Glc-NAc	349

Average *B*-factor

for all atoms (Å^2^)	71.1

r.m.s. deviations

Bond lengths (Å)	0.004

Bond angles (°)	1.03

Ramachandran plot

Most favored (%)	90.9

Allowed regions (%)	8.7

Outliers (%)	0.4

Clashscore (%)	7.7

Rotamer outliers (%)	2.4

* Values in parentheses are for the highest resolution shell (0.1Å)

\# *R*_free_ was calculated with 5% of the data
